# Engulfing to Adapt: Efferocytosis by Epithelial Cells Triggers Cell State Transitions During Tissue Remodeling

**DOI:** 10.1101/2025.07.31.668010

**Authors:** Adda-Lee Graham-Paquin, Deepak Saini, Sophie Viala, Mara KM Whitford, Mathieu Tremblay, William A Pastor, Luke McCaffrey

## Abstract

Hormone deprivation treatments are a leading approach in prostate cancer treatment. Following deprivation, the organ regresses from a fully differentiated secretory organ to one less than 50% of the original size, that has been reprogrammed to a state primed for regeneration. Despite widespread use of hormone treatments, the events linking cellular apoptotic responses following prostate hormone deprivation and cell state reprogramming are not fully understood.

We report that the absence of hormones creates a microenvironment rich in apoptotic cells that are engulfed by neighboring prostate epithelial cells. Mechanistically, we found that epithelial cells alter their metabolism and gene expression during efferocytosis. Efferocytic epithelial cells adopt a more glycolytic metabolism, resulting in elevated levels of lactate production. This correlates with increased histone Lactylation over gene regulatory elements for genes regulating autophagy and catabolism. To further demonstrate that efferocytosis is necessary to remodel tissue during regression, we expressed a mutant of the receptor protein MFGE8, MFGE8-D89E to “mask” PtdSer in the prostate epithelium and reduce efferocytosis. This effectively prevented the cell and lumen size changes hormone-deprived prostates usually go through. Our findings suggest new perspectives on how epithelial cell efferocytosis can impact tissue remodeling and altered cellular states.

## Introduction

The prostate gland is an androgen-responsive organ of the male urogenital system that secretes components of seminal fluid. The importance of androgens for prostate homeostasis is well established; testosterone depletion causes the prostate to regress to about 20% of its original weight. Conversely, restoring the androgen supply to a regressed prostate results in a full regeneration of the organ to its original size and architecture [1]. This dependence on androgens is leveraged in the treatment of prostate cancer, where chemical castration, using androgen blockers, remains a leading treatment due to its effectiveness in treating advanced prostate cancer [2]. However, incidence of recurrence is very high and often leads to a more aggressive disease [3]. This treatment plays directly on the innate dependence of the organ on androgens; much remains to be understood about how the prostate responds to androgen deprivation at both the tissue and cellular levels.

Following prostate regression, which can be triggered experimentally using androgen blockers or surgically through castration, it has been demonstrated that the epithelial cells that remain have increased stem- or progenitor potential, contributing to the regenerative potential of the organ[5-9]. One idea is that cell loss following castration could be attributed to varying degrees of dependence on androgens for survival. In this model, the ability of the prostate to regenerate can be attributed to a surviving population of resistant prostate stem cells, that are innately insensitive to androgens [7]. However, single-cell sequencing experiments as well as lineage-tracing experiments revealed that in addition to a portion of epithelial cells that continuously expressing stem-markers, all the remaining differentiated luminal cells acquire a luminal progenitor-like state in response to androgen removal [6]. This allows all residual epithelial cells to participate in regeneration once androgens have been readministered [8].

It remains unknown how differentiated epithelial cells acquire a progenitor state during regression. Through single cell ATAC sequencing experiments comparing the chromatin landscape of cells from regressed versus intact prostates, researchers saw a decrease in AR and other differentiation motifs with concurrent increase in Jun/Fos and other stem cell motifs in the open chromatin of emergent luminal progenitor population [9]. How these programs are activated remains to be elucidated.

Dying cells are present in the regressing prostate during duct remodelling, established as condensed TUNEL positive nuclei [10, 11]. Apoptotic cells are a predominant contributor to the prostatic microenvironment in the early stages of prostate regression; however, communication from these cells to their neighboring cells have not been explored. Through secretion of paracrine factors, apoptotic cells can induce death or proliferation in surrounding cells, ultimately regulating tissue growth in the area [12].

One interaction that exists between the apoptotic cell, and its phagocyte, is that of phosphatidylserine exposure on the plasma membrane of the apoptotic cell to initiate their engulfment [13]. Macrophages, the most well characterized phagocytic cells, have been known to alter their gene expression, metabolism, and behavior within a tissue with exposure to apoptotic cells [14-17]. Here we demonstrate that the phagocytosis of apoptotic cells by epithelial cells changes their metabolic and differentiation programs during prostate regression, ultimately contributing to the morphological changes of epithelial cells in the prostate.

## Results and Discussion

### Apoptotic cell clearance in the regressing prostate is due to efferocytosis by epithelial cells

To understand cellular events following androgen-deprivation, we performed surgical castration in mice and followed the effect on prostate tissue over time (Figure 1A). As expected, the prostate regressed to less than half of its original size (Figure 1B, Supplemental Figure 1A). Following prostate regression, we observed that epithelial cells reduced their size, the duct lumen volume decreased, and ducts displayed less epithelial folding (Figure 1C, Supplemental Figure S1B-D.). In addition to these morphological changes in response to androgen-deprivation, studies over the past several decades reported there is massive cell loss [18] due to a surge in apoptotic cell death in the prostate [10, 19]. This characterization has mostly been limited to the ventral lobe of the prostate [10, 11, 19-22]. We observed that the degree and timing of peak apoptosis in the four lobes of the prostate are different, with the lateral lobe experiencing lower levels of apoptosis, despite all lobes expressing similar levels of androgen receptor [23] (Figure 1D-F). A large portion of cell loss may also be attributed to desquamation, observed first by Rosa-Ribeiro et al [24]. where large strips of epithelial cell sheets shed from the ducts. By inspecting this process further, we observed that desquamation occurred at low levels in steady state adult prostate and was more prevalent in early response to androgen deprivation, reducing to sub-homeostatic levels by 4 days after castration (Figure 1G-I, Supplemental Figure 1G-I). The timing of this early response is consistent with a direct response to the sharp drop in androgens following castration. Programmed cell death, through cleaved caspase 3 (cl-cas3) dependent apoptosis, occurs concurrently with desquamation, with no correlation between the levels of the two events within a given duct (Figure 1I, Supplemental Figure 1J).

**Figure 1.**
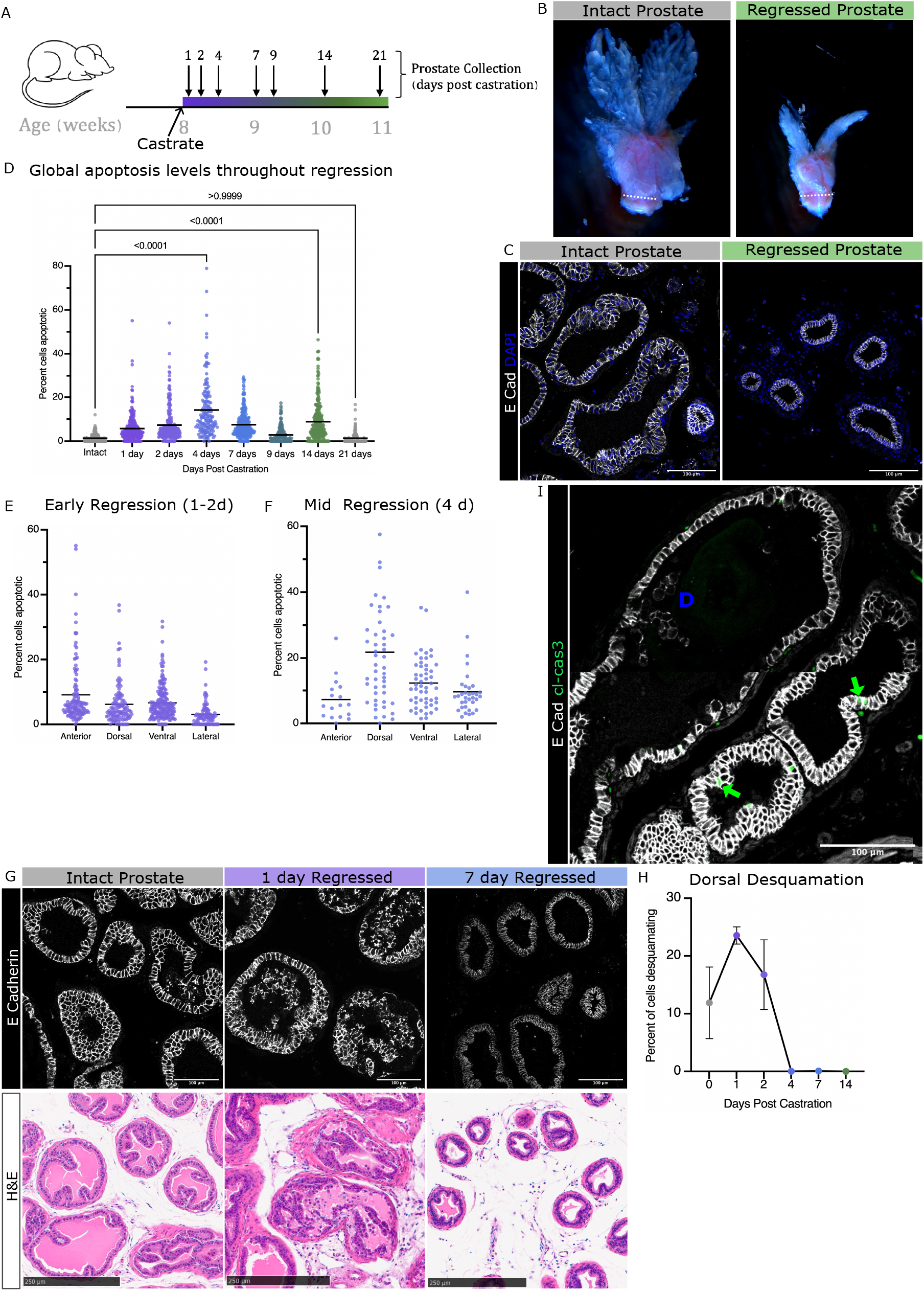
Morphological regression of the mouse prostate is due to desquamation, apoptosis and changes to cell shape. A. Schematic of timeline for castration and prostate collection throughout regression. B. Representative images of intact and regressed prostates. Dotted line to show urethral width, which is unchanged in regression (average width 1.39mm). C. Tissue sections, representative of dorsal lobe ductal and cellular morphology of intact and 21-day regressed prostates, immunofluorescence stain for E Cadherin to show epithelial membranes. D. Scatter plot showing global apoptosis levels throughout prostate regression. Each data point represents the percent apoptotic (TUNEL+) nuclei per duct, compiled from 3 mice per timepoint, from 50-75 ducts per mouse. ANOVA statistical test between intact_1d p<0.0001, intact_2d p<0.0001, intact_4d p<0.0001, intact_7d p<0.0001, intact_9d p=0.1086, intact_14d p<0.0001, intact_21d p>0.999 E. Apoptosis levels per lobe during early regression (1-2 days post castration). Each data point represents the percent apoptotic (TUNEL+) nuclei per duct, compiled from 3 mice per timepoint, from 8-10 ducts per mouse per lobe. F. Global apoptosis levels per lobe mid-regression (4 days post castration). Each data point represents the percent apoptotic (TUNEL+) nuclei per duct, compiled from 3 mice per timepoint, from 8-10 ducts per mouse per lobe. G. Immunofluorescent (TOP) and H&E (BOTTOM) images of desquamation in the dorsal lobe of prostates from Intact, 1 and 7 days after castration. H. Average percent of detached cells in the duct in the dorsal lobe over time, Error bars= SEM. I. Tissue section of a duct with immunofluorescence stains for E-cadherin and cleaved caspase 3 in which desquamation (D) and apoptosis (Green arrows) can be seen concurrently.

Since the majority of apoptotic cells were observed within the epithelium but rarely in the lumen (Supplemental Figure 2A), a method to clear them from the epithelium is required. To identify the predominant cell clearance mechanisms of these apoptotic cells, we performed rigorous assessment of tissue sections of the regressing prostate using multiplexed immunofluorescent staining with markers for apoptotic cells (cleaved-caspase 3), DNA fragmentation (TUNEL), macrophages (CD68), and epithelial cells (E-cadherin). Macrophages, which are canonically responsible for dead cell clearance, are scarce, but when found were restricted within the stromal compartment and contained markers of dead/dying cells (Supplemental Figure 2B-D). Strikingly, we observed extensive engulfment of apoptotic cells by epithelial cells during prostate regression. As discerned by cleaved-caspase 3 (cl-cas3) and TUNEL staining, apoptotic cells were found at different stages of degradation within epithelial cells and were contained within LAMP1+ phagolysosome bodies (Figure 2A-C). As apoptosis levels rise in response to androgen-deprivation, so too does the prevalence of epithelial cell engulfment by neighboring epithelial cells, making it the predominant mechanism of dead cell clearance in the regressing prostate (Figure 2D, Supplemental Figure 2E,F). Unlike macrophages, which migrate through tissues collecting corpses, these epithelial phagocytes seem to “share the load” as 10% of epithelial cells in a duct can be seen actively degrading apoptotic cells at a given moment during regression (Figure 2E). We compared the waves of efferocytosis and their timing between lobes, and we observed that the presence of early stages (apoptotic cells not yet engulfed) is followed by the peak in active engulfment (cl-cas3 positive bodies within epithelial cells), followed then by peaks in late engulfment (cl-cas3 negative, TUNEL positive bodies or debris within epithelial cells). However, these waves occur earlier in the dorsal and anterior lobes compared to ventral and lateral, with peaks in efferocytosis occurring by 4 days after castration and a smaller surge again after 14 days (Figure Supplemental Figure 2G-J). We therefore focused our analysis on dorsal and anterior lobes to control for this difference in timing. To see if the expression levels of genes involved in phagocytosis change to accommodate the waves of cell death during regression, we reanalyzed the single cell sequencing dataset generated by Karthaus et al.[8], looking specifically at epithelial cells. Several genes essential for the internalization of apoptotic bodies like Rac1 and Rab7 show increased expression in the epithelium during the surge in apoptosis seen 14 days post-castration (Figure 2F).

**Figure 2.**
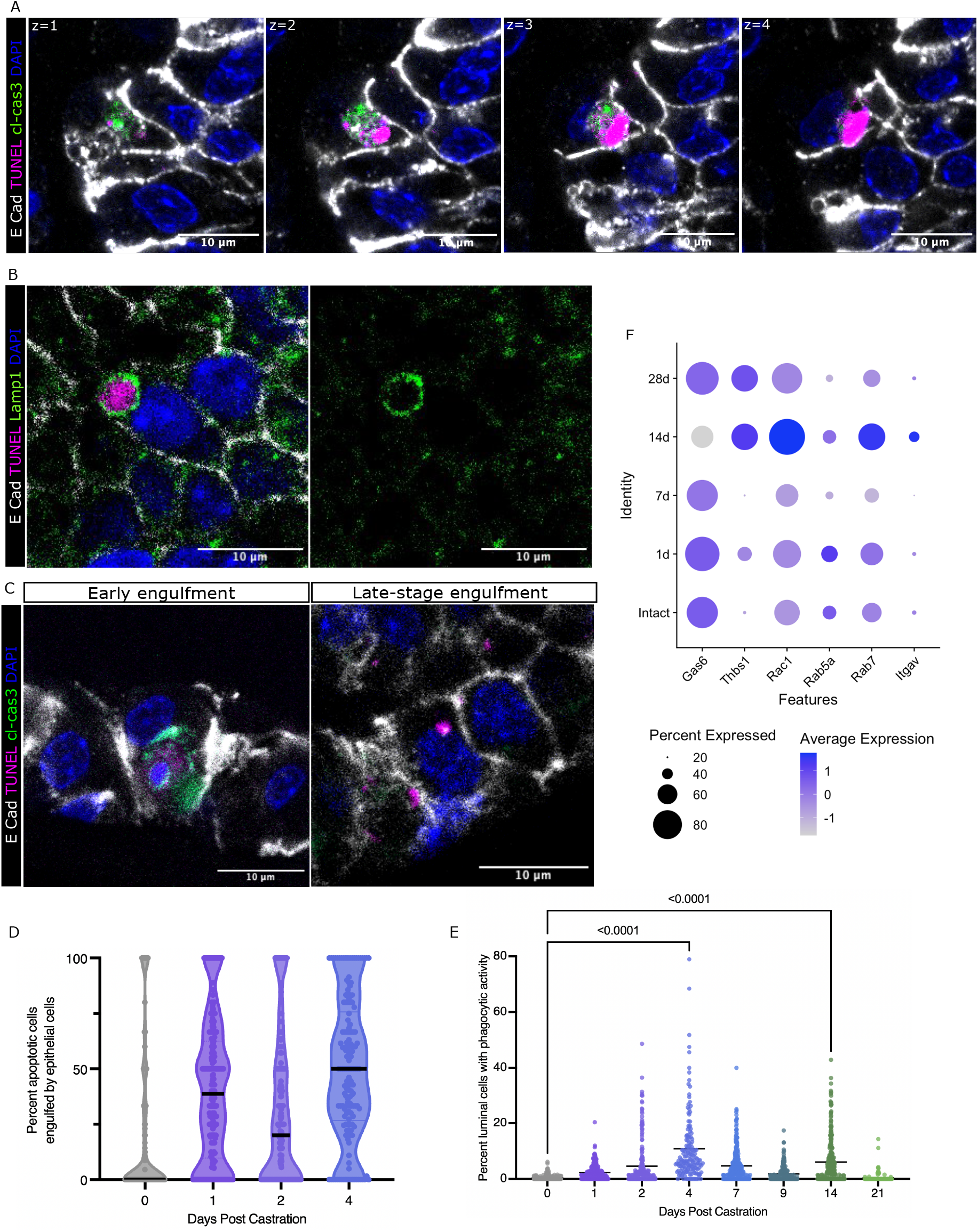
Epithelial cells in the prostate are phagocytic in response to castration-induced apoptosis. A. Z-stack (1 m increments from z=1 to 4) of immunofluorescent stained tissue section of 1-day regressed prostate showing an epithelial cell with an apoptotic cell inside of it. E Cadherin marks the membranes, the apoptotic cell within has TUNEL positive signal to indicate fragmented DNA and cl-cas3 positive cytoplasmic components. The efferocytic cell’s healthy nucleus can be seen as TUNEL negative DAPI in stack z=3,4. B. Immunofluorescence-stained tissue section of 4 day regressed prostate showing a degrading apoptotic body (identified by TUNEL +ve foci) within a phagolysosomal body (LAMP1+ve vesicle) of healthy epithelial cell. C. Immunofluorescence-stained tissue sections of 1- and 4-day regressed prostates to show early and late stages of cell engulfment. All dying cell components can be visualized by TUNEL and cl-cas3 foci. D. Violin plots showing the increase in epithelial efferocytosis in early regression compared to intact prostates. Each data point represents the percentage of total apoptotic cells in or around a duct that can be found within an epithelial cell. Solid black line represents the median. Dotted lines show the other quartiles. Data compiled from 3 mice per timepoint, from 50-75 ducts per mouse. Mean values (not shown on graph) are as follows: intact 18.79%, 1d 39.84%, 2d 30.21% and 4d 49.53%. E. Scatter plot of the phagocytic activity of luminal cells throughout regression. Each data point represents the percent of cells within a duct containing an apoptotic body, compiled from 3 mice per timepoint, from 50-75 ducts per mouse. Bar indicates the mean. ANOVA statistical test between intact_1d p<0.0001, intact_2d p=0.0034, intact_4d p<0.0001, intact_7d p<0.0001, intact_9d p=0.1089, intact_14d p<0.0001, intact_21d p>0.999. F. Dot plot of single cell gene expression of various phagocytic components throughout regression, in epithelial cells.

### Prostate epithelial cells adapt their metabolism to accommodate efferocytic demand

To understand the metabolic change in epithelial cells following castration, we performed LC/MS quantification of metabolites. We specifically chose to examine metabolic change in the prostate during regression 4 days after castration since it corresponds to the most active period of active cell death and engulfment and, compared it to both intact and fully regressed prostates (21 days post-castration). Unbiased analysis of KEGG metabolic pathways enriched only at 4 days after castration revealed that nucleotide metabolism (uracil and uric acid) and amino acid catabolism (arginosuccinate) were most significantly increased (Supplemental Figure 3A). We propose that it is due to the increased catabolic load on the efferocytes, having to degrade their apoptotic neighbors (Figure 3A-C, Supplemental Figure 3A). Interestingly active regression (4d regressed) is associated with higher global NAD levels, increased NAD/NADH ratio and elevated creatine phosphate. This is consistent with an enhanced energetic demand and capacity needed by epithelial cells to perform this process (Figure 3D-F). In line with increased efferocytic activity and lysosomal targeting, a transient increase in Mannose-6-phosphate levels was observed during regression (Supplemental Figure 3B).

**Figure 3.**
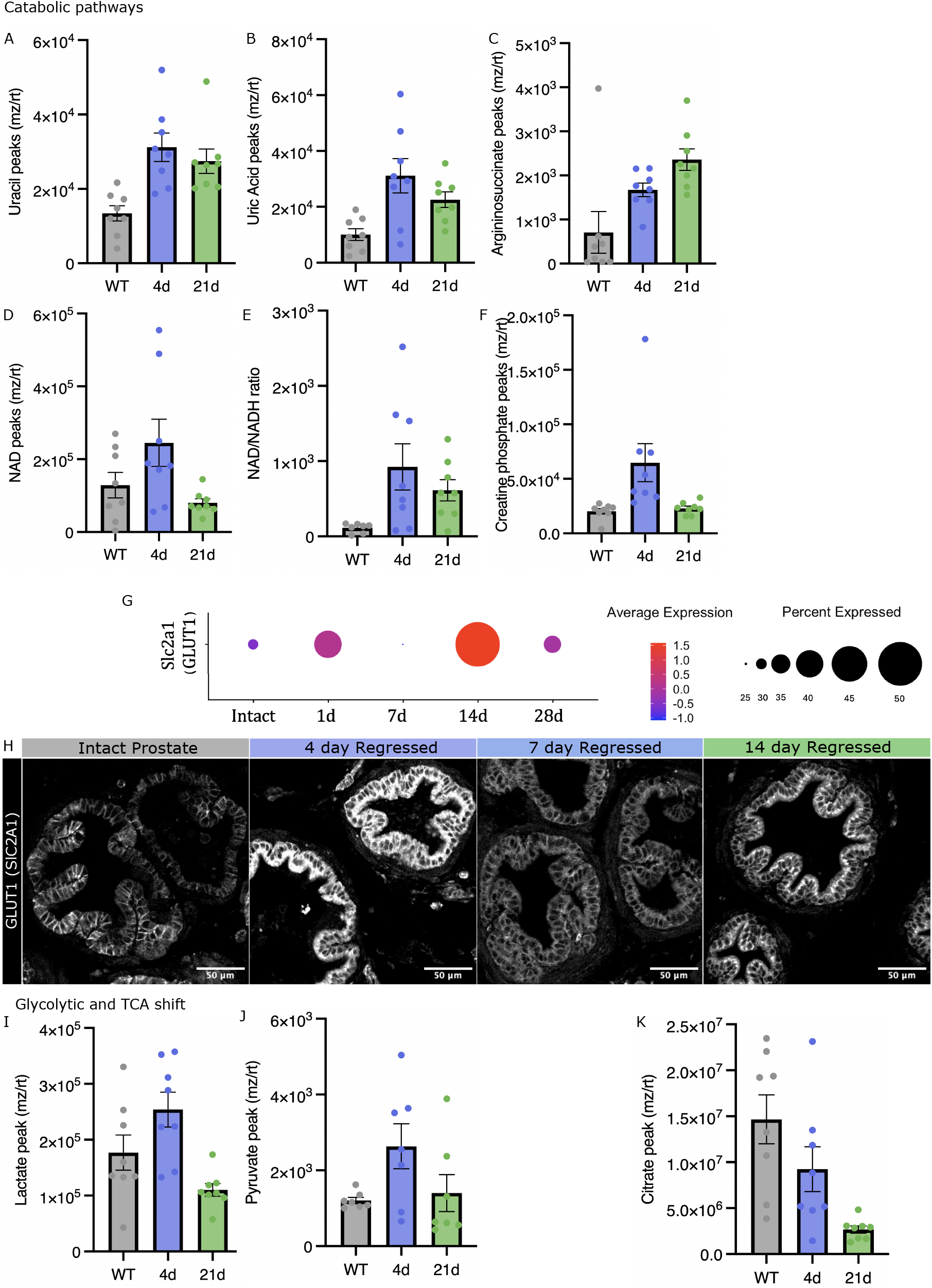
Prostate epithelial cells adapt their metabolism to accommodate efferocytic demand. A-F. Levels of metabolites present in anterior & dorsal lobes of intact, 4- and 21-regressed prostates. Each data point represents the metabolite level per mouse, 8 mice per condition. A. Uracil between Intact_4d p=0.0018, 4d_21d p=0.6828, Intact_21d p=0.0126. B. Uric Acid between Intact_4d p=0.0042, 4d_21d p=0.3212, Intact_21d p=0.1020. C. Arginosuccinate between Intact_4d p=0.1067, 4d_21d p=0.3047, Intact_21d p=0.0041. D. NAD between Intact_4d p=0.1616, 4d_21d p=0.0339, Intact_21d p=0.7069. E. NAD/NADH ratios between Intact_4d p=0.0314, 4d_21d p=0.5305, Intact_21d p=0.2296. F. Creatine phosphate between Intact_4d p=0.0148, 4d_21d p=0.0226, Intact_21d p=0.9798. G. Dot plot of single cell gene expression of Slc2a1 (GLUT1) throughout regression, in epithelial cells of the anterior lobe. H. Tissue sections from intact, 4-, 7- and 14-day regressed prostates immunofluorescence-stained for GLUT-1 I-K. Levels of metabolites present in anterior & dorsal lobes of intact, 4- and 21-regressed prostates. Each data point represents the metabolite level per mouse, 8 mice per condition. I. Lactate between Intact_4d p=0.1208, 4d_21d p=0.0.00125, Intact_21d p=0.1976. J. Pyruvate between Intact_4d p=0.0.0872, 4d_21d p=0.1523, Intact_21d p=0.9488. K. Citrate between Intact_4d p=0.1834, 4d_21d p=0.0914, Intact_21d p=0.0016. Error bars represent SEM. ANOVA statistical tests for differences.

Macrophages have been observed to undergo many changes in response to apoptotic cell exposure, often attributed to a state of “polarization”, a term used to describe function and behavior of the cell. We therefore explored the possibility that phagocytic epithelial cells also alter their cell state in response to the engulfment of apoptotic cells. One amazing influence that apoptotic cells have on macrophages is to directly alter their gene expression of solute carriers (SLC) transporters, resulting in a change in their metabolic activity [15]. In macrophages, Slc2a1 (GLUT1), which encodes a glucose transporter is upregulated in response to apoptotic cell docking, is proposed to be essential in driving glycolysis-fueled efferocytosis. Temporal mRNA expression in the anterior prostate of the GLUT1 transporter coincided with peaks of apoptotic cell engulfment within the lobe (Figure 3G). Indeed, through immunofluorescent staining of GLUT1, we see an increase in protein expression in 4d and 14d regressed prostates, coinciding with the two waves of apoptosis observed during regression (Figure 3H). In addition, a slight increase in the amounts of pyruvate and lactate were detected during efferocytosis, followed by a decrease in the fully regressed prostate (Figure 3I,J). Also, there was downshift in levels of TCA metabolites, citrate, malate, fumarate and succinate, indicative of a push towards aerobic glycolysis during active efferocytosis (Figure 3K, Supplemental Figure 3C-H).

### Histone lactylation marks in early regression align with genes important for efferocytosis and autophagy

Given the metabolic reprogramming in the epithelial phagocytic cells in the prostate in response to castration, the effect was analysed over time to see if it could have any lasting on the phagocytic epithelial cells. Lactate produced by glycolytic cells has been shown to modify cell states through enhanced nuclear protein lactylation and epigenetic regulation by histone lysine-lactylation (KLa)[16, 25]. We first determined if there were changes in nuclear lysine-lactylation following castration at timepoints corresponding to early (1d) and mid-efferocytosis (4d). Through immunofluorescent imaging of tissue sections, we observed some nuclear lysine-lactylation at 1d regressed, which was increased by 4d regressed (Figure 4A). Since histones have been shown to be a protein that can be lactylated [14, 23], we looked for lysine-lactylation between intact and regressed prostate. Interestingly, we found an abundance of lysine-lactylation at 15kDa, potentially corresponding with Histone H3 which presents a similar band at 15kDa on gel (Supplemental Figure 4A,B).

**Figure 4.**
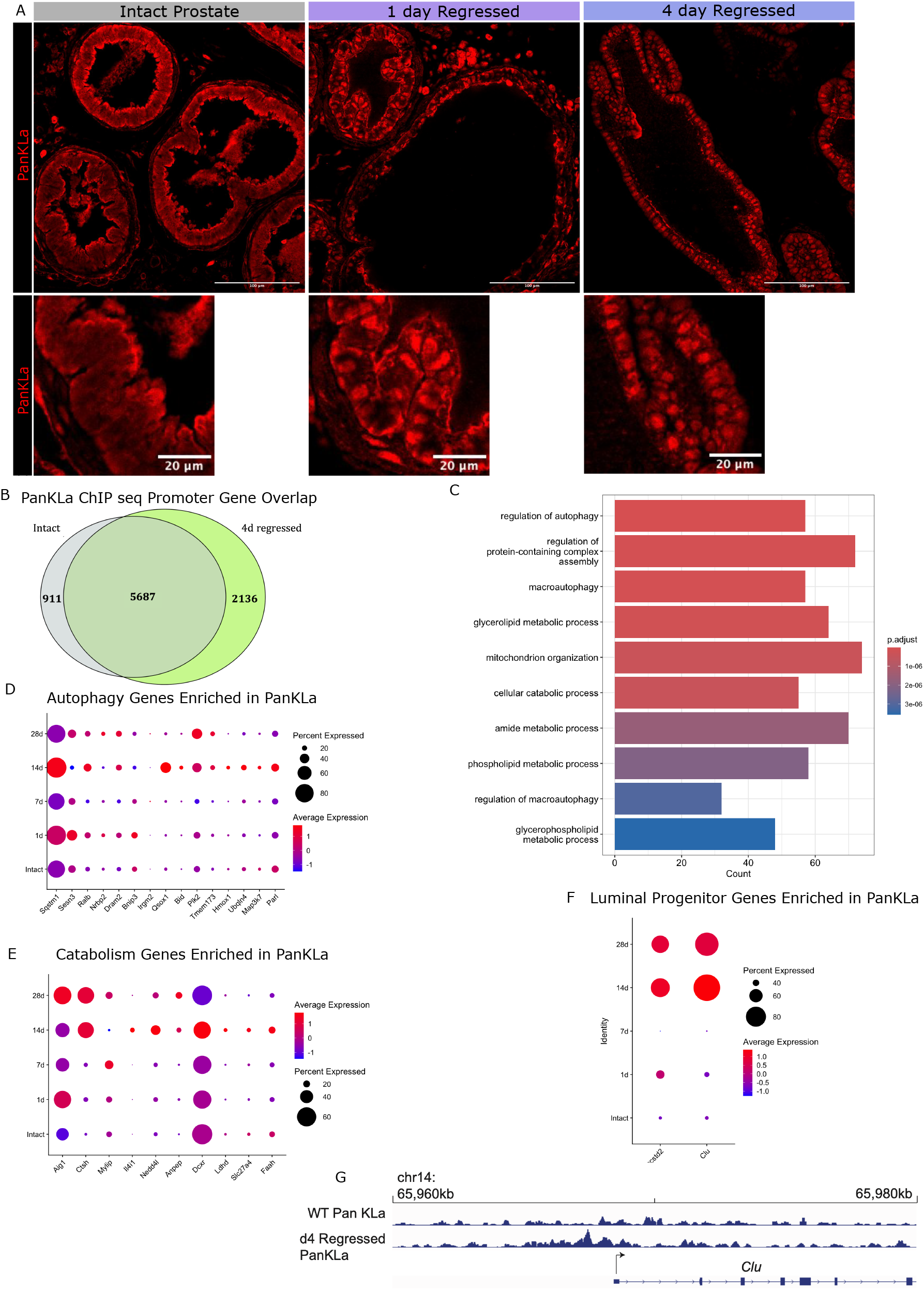
Histone lactylation marks in early regression align with genes important for efferocytosis and autophagy. A. Tissue sections from intact, 1-, 4-day regressed prostates immunofluorescence-stained for Lysine Lactylation (PanKLa) B. Venn diagram of genes with promoter PanKLa peaks in intact versus 4-day regressed prostates C. GO terms of enriched pathways for genes with promoter lactylation exclusively in 4-day regressed prostates D. Dot plot of single cell gene expression of autophagy genes with promoter lactylation exclusively in 4-day regressed prostates throughout regression, in epithelial cells. E. Dot plot of single cell gene expression of catabolism-related genes with promoter lactylation exclusively in 4-day regressed prostates throughout regression, in epithelial cells. F. Dot plot of single cell gene expression of luminal progenitor markers with promoter lactylation exclusively in 4-day regressed prostates throughout regression, in epithelial cells. G. Example track of promoter lactylation (PanKLa ChIP seq) upstream of the gene Clusterin (Clu).

To explore if nuclear lysine-lactylation has a functional outcome, we performed ChIPseq on the dorsal and anterior lobes of intact and actively regressing prostates (4d regressed) and mapped the lysine-lactylation peaks to the mouse genome. The majority of lysine-lactylation peaks associates with gene promoter regions (<=1kb upstream of genes), representing 60.17% of in intact and 50.76% in regressing prostates (Supplemental Figure 4C,D). While 5687 genes were found lactylated in both conditions, we noted that 2136 genes had exclusively promoter lactylation in regressing prostates, while 911 genes were exclusively lactylated in intact prostates (Figure 4B). Genes targeted by lactylation upon regression showed an enrichment in genes related to autophagy, regulation of apoptosis, and catabolic processing when performing Gene Ontology (GO) enrichment (Figure 4C). Using a publicly available dataset, we found that several of these genes were transcriptionally increased in either early or late regression when compared to the intact prostate (Figure 4D,E, Supplemental Figure 4E,F). Interestingly, we found that the promoter of *Clu* and *Tacstd2*, genes previously associated with the luminal cell state transition, present high lysine-lactylated levels in regressing prostates[8, 9] (Figure 4F, Supplemental Figure 4G). Although we cannot confirm a functional role of these lactylation marks in the regressing prostate, their differential presence on genes known to change expression during regression presents an intriguing starting point for future investigation.

### Impairing efferocytosis impedes prostate regression

To investigate the degree to which efferocytosis influences prostate regression and the associated cell state transition, we developed a mouse model that specifically impede cell engulfment without blocking apoptosis itself throughout the regression process. To do this, we generated an inducible MFGE8-D89E allele, which conditionally express the dominant negative form of MFGE8 (D89E mutation)(Figure 5A, Supplemental Figure 5A,B). This mutant has been shown to efficiently block phagocytosis by both macrophages and epithelial efferocytes in *ex vivo* culture systems [26, 27], by masking phosphatidyl serine (PtdSer) on apoptotic cells. Using inducible K8CreERT2 allele to target specifically luminal cells (Figure 5A), activation of the Cre allele with tamoxifen induced MFGE8-D89E expression a week prior to castration hindered prostate regression. By 21 days post castration, MFGE8-D89E expressing prostates had undergone only a 30% decrease in size when regressed wildtype prostates by this time decrease by 56% after castration (Figure 5B-D, Supplemental Figure 5C-E). Furthermore, changes in cell and duct geometries associated with regression (Supplemental Figure 1A-D), were impeded by MFGE8-D89E expression. The cell height remained larger, and the lumen size was not as reduced as was normally observed (Figure 5E, F, Supplemental Figure 5F-H). In addition, there were more apoptotic cells remaining in the tissue, likely resulting from a reduce rate of efferocytic process when PtdSer are masked (Figure 5G, Supplemental Figure 6A). To see if there was an effect on the acquired luminal progenitor state in the regressed prostate, we explored the expression of one of the key markers of this state, Tacstd2 (TROP2) by immunofluorescence and saw a failure of the cells to increase their membrane Tacstd2 levels to what is normally observed by 21 days post castration (Figure 5H,I). Our results show that blocking efferocytosis affect the tissue regression process and potentially influence the transformation into luminal progenitors.

**Figure 5.**
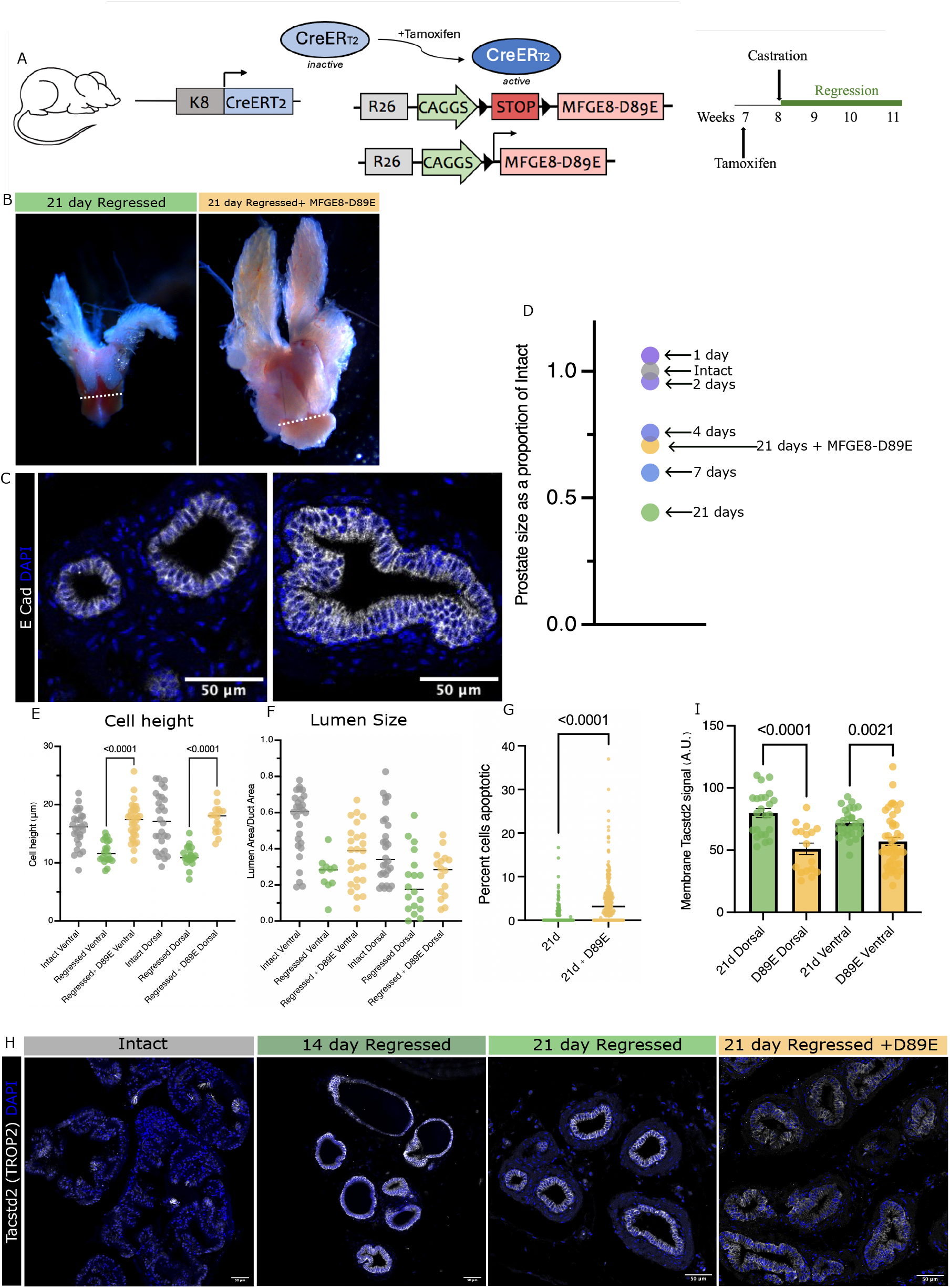
Impairing efferocytosis impedes prostate regression. A. Schematic of mouse model for impaired efferocytosis during prostate regression B. Whole mount images of regressed prostates 21 days after castration with and without MFGE8-D89E induction to impair efferocytosis. Dotted line to show urethral width, which is unchanged in regression (average width 1.39mm). C. Tissue sections, representative of dorsal lobe ductal and cellular morphology of 21-day regressed prostates with and without impaired efferocytosis, immunofluorescence stain for E Cadherin to show epithelial membranes. D. Average prostate size as a proportion of average intact prostate size at 1-, 2-, 4-, 7-and 21-days post-castration, with the average size of 21 days +MFGE8-D89E prostates labeled in comparison. E. Cell height, each data point represents average cell height per duct. N=3 mice, 10 ducts per mouse per lobe F. Lumen size, quantified as ratio of lumen area/ ductal area to control for changing duct size with regression, each data point represents one duct. N=3 mice, 10 ducts per mouse per lobe. G. Scatter plot showing apoptosis levels at 21 days post-castration with and without MFGE8-D89E. Each data point represents the percent apoptotic (TUNEL+) nuclei per duct, compiled from 3 mice per timepoint, from 8-10 ducts per mouse. H. Tissue sections from intact, 14-, 21- and 21-day with MFGE8-D89E regressed prostates immunofluorescence-stained for Tacstd2 and DAPI. I. Quantification of membrane intensity of Tacstd2 in dorsal and ventral lobes. Each data point represents the membrane intensity from a single duct, compiled from 3 mice per timepoint, 10 ducts per mouse.

### Efferocytosis-Induced Reprogramming Promotes Epithelial Plasticity in the Regressing Prostate

Apoptotic cell signaling has long been known to contribute to tissue reorganization in instances of wound repair [28, 29] and in tissue regeneration in both amphibians [30], and mammalian tissues [31]. The role of macrophages, or “professional” immune phagocytes in both dead cell clearance and tissue remodeling has also been well established [32]. Here, we demonstrate that efferocytosis by non-immune cells, a phenomenon that has been observed in many tissues and organisms [33], can also influence tissue remodeling as well as epithelial cell state.

By blocking efferocytosis throughout the regression process using MFGE8-D89E, we were able to definitively demonstrate a role for efferocytosis in this process. Notably, the prostate regression is not fully prevented when efferocytosis is impeded. This regression can be attributed to the initial cell loss through desquamation which does not involve efferocytosis, although absolute cell numbers and total contribution of desquamation to organ regression have not been quantified.

It must be understood that as MFGE8-D89E is a secreted factor that “masks” dying cells, cell engulfment and signaling through PtdSer to the macrophages in the surrounding tissue is also obstructed in our model, and therefore we cannot ignore the potential effect of this. Macrophages are unquestionably also involved in clearing dying cells in the regressing prostate [21, 34]. However, the contribution of the epithelial cells in taking on this apoptotic cell load is greater during prostate regression.

Using our understanding of macrophage biology, through metabolic reprogramming in response to efferocytosis and the resulting effect on transcriptional networks that influence macrophage behaviour [17], we suggest a model by which epithelial cells acting as phagocytes undergo a similar experience. Not only do epithelial cells respond by becoming more glycolytic, but the accumulated lactate resulting from this metabolic switch leads to an epigenetic reprogramming. These lactylation marks differentially distribute across the genome in the regressing prostate could help these cells adapt to their new phagocytic role, by stimulating transcription of genes essential for autophagy and catabolism of various metabolites ingested from dying cells. Interestingly it has been shown in both migrating neural crest cells and activated macrophages that lactylation marks acquired through a similar mechanism (increased lactate due to glycolytic enhancement) correlates with areas of open chromatin and actively transcribed genes [16, 25]. In addition, the gene networks activated by lactylation in these systems were also associated with cell plasticity. Therefore, the role of lactylated gene networks and epithelial cell plasticity must be further explored. It has been increasingly evident that androgen deprivation itself triggers a response in epithelial cells to acquire a luminal progenitor state, possibly explaining the actuality of aggressive castration resistant prostate cancer [35]. Here we establish a role for apoptotic cell signaling, specifically cell engulfment, in the transformation of both tissue and cell during prostate regression following androgen-deprivation. This provides a deeper understanding into the potential mechanism for these cells becoming progenitors, and potentially for prostate cancer cell response to androgen deprivation. Based on our results in normal prostate, exploring the role of efferocytosis in prostate cancer response to androgen deprivation would be an interesting avenue of research, as it could lead to improved treatment options.

## Methods

### Animal Studies

#### Ethics Statement

All experimental procedures performed with animals were conducted in compliance with the Canadian Council of Animal Care (CACC) requirements for research using mice and overseen and approved by the McGill University Animal Care Committee.

Mice were housed in the Goodman Cancer Institute Animal Facility, with a 12 h cycle of light and darkness, a temperature of 20–24 °C, and a relative humidity of 45–65%.

#### Experimental mice

All experimental mice were kept in a C57/BL6 background. Wildtype C57/BL6 mice were obtained from Charles River Laboratories and acclimated to the McGill Animal Facility for at least one week prior to any experimental procedures.

K8Cre^ERT2^; MFGE8-D89E mice were generated through in-house breeding between K8Cre^ERT2^ [6] mice, described previously, and MFGE8-D89E (Rosa26-pCMV-pβactin-LSL-MFGE8-D89E-WPRE) mice.

MFGE8-D89E mice were generated through CRISPR-Cas9 homology-directed-repair gene editing, by targeted introduction of a pCMV-pβactin-LSL-MFGE8-D89E-WPRE fragment to the Rosa26 locus using sgRNA guide (CCA GTCTTTCTAGAAGATGGGCG). Our donor template was generated by clonal modifications of the Ai9 Vector (addgene Plasmid #22799), previously used to create the tdTomato mouse line [36], a very strongly expressing fluorescent reporter line. This was done by swapping out the tdTomato sequence for MFGE8-D89E [26] and removal of unnecessary NeoR/KanR sequences. The MFGE8-D89E-FLAG construct was kindly given to us by Dr. Shigekazu Nagata. Genotyping was performed by PCR amplification using primers 5′-gctgatccggaacccttaat-3′ and 5′-TGGCAGATGTATTCGGTGAA-3′.

Unless explicitly stated otherwise, all experiments were done using 8–10-week-old adult mice. Tamoxifen was administered to all experimental and control mice through intraperitoneal injection of tamoxifen dissolved in sunflower oil. Injection of 3mg tamoxifen was given 3 times, over the span of 5 days (9mg total), as this dosage was found previously to lead to full Cre recombination of the stop cassette located in the Rosa26 locus of tdTomato mice [37, 38]. Castrations of adult mice were performed on adult mice in accordance with McGill SOP207), and prostate tissue was collected at 24hr, 48hr, 4-day, 7-day, 9-day, 14-day or 21-day time points after castration, as indicated, and used for subsequent experiments.

#### Tissue isolation and handling

Tissues were dissected in cold sterile PBS. Those prostates used to generate tissue sections for histological analysis were fixed in 4% PFA at 4°C for 24 hours and processed for paraffin embedding, where 4μm sections were collected. All prostates used for biochemical analyses were processed from fresh tissue as described.

#### Immunofluorescent Staining

Sectioned tissue embedded in paraffin were warmed to 56°C and then deparaffinized in Histoclear (National Diagnostics Cat# 50-899-90147) for 5 min, then again for 3 min, then slowly rehydrated by 3-minute increments in 100%, 95% 70% ethanol, followed by a slow flow-over of dH2O to bring the slides into 100% water. Following a PBS rinse, antigen retrieval was done in Tris-EDTA antigen retrieval buffer, in a pressure cooker for 7 minutes, followed by gradual cooling and another PBS rinse. All slides were then blocked in SuperBlock Blocking Buffer (Thermofisher CAT# 37515) at room temperature for 1 hour. Primary antibodies were diluted in PBS and slides were incubated for 48-72hr at 4°C. Following four 5-minute washes in PBS, PBS, PBS, PBST respectively, slides were incubated in PBST + 2% Normal Donkey Serum + 0.2% Triton-X-100 + respective secondary antibodies (all diluted at 1:500) at room temperature for 1 hour. In the case of slides stained using TUNEL (In situ Death Detection Kit; ROCHE Cat# 11684795910), secondary antibodies were diluted at 1:500 and the enzyme was used at 1:15 in the TUNEL buffer provided; these slides were incubated for 45 minutes at 37°C. Following three 10-minute washes in PBS, PBST, PBS respectively, slides were mounted with Mowiol mounting media.

#### Imaging and analysis

Whole mount images of prostate organs were taken using the Zeiss Lumar V12 stereomicroscope at 6.4X magnification. Areas of the lobes were then measured from 2D images of all prostates in the same orientation (normal for anterior, side for dorsal/lateral and ventral).

Slides stained with H&E were scanned on Nanozoomer S210 Brightfield scanner at 20X magnification. Fluorescence images were acquired using either a Zeiss LSM800 or LSM-P-PMT confocal laser scanning microscope, equipped with a 20x/NA = 0.80 dry plan apochromat objective, from the McGill University Advanced BioImaging Facility (ABIF, RRID:SCR_017697).

Image analyses were performed with ImageJ v1.54p. Desquamating cells were counted as E-Cadherin positive cells found within the lumen, no longer connected to the epithelial layer anchored to the basal membrane. Apoptotic events were quantified as the number of cl-cas3 positive or TUNEL positive foci located within the duct. Apoptotic cells were either classified as being within the epithelial compartment (marked by E-Cadherin), within the lumen, or within the ducts stromal compartment (located just outside epithelium basally). All “early” apoptotic events encompass cl-cas3 positive cells not yet being engulfed. All “mid” engulfment events encompass cl-cas3 positive bodies (1.5μm or larger) found within an epithelial cell. All “late” engulfment events encompass TUNEL positive, cl-cas3 negative bodies (0.8μm or larger) found within an epithelial cell. Cell height per duct was measured by averaging the (apical to basal) length of 5 cells per duct. Lumen and Ductal areas were measured manually in Image J. Tacstd2 membrane signal quantification was done using image j and measuring the signal intensity of the membrane using E-Cadherin as membrane marker. (Script available upon request).

## Statistics

Statistics were performed using Prism 10.0 software (Graphpad). Comparisons of two conditions were done using unpaired two-tailed Student’s *t* test, and statistical comparisons of three or more conditions were done using one-way ANOVA.

### Metabolite measurements and analysis: LC/MS

Prostates from 24 mice (8 intact, 8-4 days after castration and 8-21 days after castration) were fine dissected and anterior and dorsal lobes were flash frozen in liquid nitrogen and crushed using CP02 Automated Dry Pulverizer. Samples were stored at −80C until used. 5mg of tissue per sample were weighed out and metabolite extraction was done as follows. Chilled sample was resuspended in 50% MeOH with 1mg/ml N-Ethylmaleimide (NEM). Acetonitrile (ACN) was added to a total of 36% vol and samples were homogenized using a bead beater and 2.8mm beads for 2 times 2 minutes at 30Hz. Dichloromethane was added to 50% total volume and ice-cold molecular grade H_2_O to 25% total volume. After a 1-minute vortex, layers partitioned for 10 minutes on ice, then spun at (1500 x g) for 10 minutes at 1°C. The aqueous phase was collected and after being dried in a SpeedVac, were resuspended in injection buffer to be injected into the Agilent LC/MS UPLC/QQQ spectrometer and analyzed using Masshunter software (Agilent Technologies). Metabolites were identified by comparison of the ion features in the experimental samples to a reference library of chemical standard entries that included retention time and molecular weight (*m*/*z*). All peaks were verified manually for accuracy. Peak intensities were quantified using area-under-the-curve (AUC) for all samples. Differential analysis was done using Metaboanalyst after autoscaling data (mean-centered and divided by the standard deviation of each variable) and all significantly different metabolites between intact and 4 day post castrated samples were submitted through KEGG pathway analysis using Metaboanalyst.

### Western Blot

Protein extracts were extracted from flash frozen tissue ground with mortar and pestle, using RIPA lysis buffer (10 mM Tris–HCl (pH 8.0), 140 mM NaCl, 1% Triton X-100, 0.1% sodium dodecyl sulphate, 0.1% deoxycholate, 1 mM EDTA) supplemented with protease and phosphatase inhibitor cocktails (Roche 11697498001). After being run for 2 hours at 100V on a 12% resolving polyacrylamide gel, proteins (20ug/sample) were transferred to an Immobilon FL PVDF membrane (Millipore IPFL00010) and probed with the following antibodies diluted in Odyssey blocking buffer (LI-COR): anti-PanKLa, and anti-Histone H3. Rabbit IR dye secondary antibodies (LI-COR Biosciences) were used at 1:20 000. Blots were scanned with the Odyssey imaging system (LI-COR).

### Chromatin Immunoprecipitation

Anterior and dorsal lobes collected from 4 wildtype prostates and 4 prostates collected 4 days after castration, like-conditions were pooled. The prostates were minced with scissors and then and digested into single cells using collagenase hyluronase (Stemcell Technologies), for 1 hour at 37C, followed by 5-minute incubation in 2X TrypLE (Gibco) and rinse with PBS. These single cells were then crosslinked in 1% paraformaldehyde for 10 min and quenched with the addition glycine to final concentration of 0.14M for 10 min to stop crosslinking. Cells were lysed in lysis buffer (Triton buffer for 15 min (10 mM Tris-HCl, pH 8.0, 10 mM EDTA, 0.5 mM EGTA, 0.25% Triton X-100) followed by NaCl buffer for 15 min (10 mM Tris-HCl, pH 8.0, 1 mM EDTA, 0.5 mM EGTA, 200 mM NaCl)) and moved to a modified RIPA buffer (10 mM Tris-HCl, pH 8.0, 10 mM EDTA, 0.5 mM EGTA, 0.1% SDS) and sonicated with the M220 ultrasonicator (Covaris) for 5 minutes total (6*30s bursts) at 200 cycles per burst, 20% duty factor, peak intensity power of 75, at 7°C. After pre-clearing the protein lysate using magnetic Protein A/G beads (Millipore #16-663) for 1 hour, the lysate was transferred to a new tube where either αPanKLa (PTM-1401RM) or αH3K4me3 (CST CD42D8) were added and incubated overnight with rotation. DNA/Protein-bound antibodies were then bound by washed magnetic beads through 2-hour incubation at 4°C.

Bound beads were then subject to multiple washes with the following buffers A: 50 mM HEPES, 1% Triton X-100, 0.1% deoxycholate, 1 mM EDTA, 140 mM NaCl, B: 50 mM HEPES, 0.1% SDS, 1% Triton X-100, 0.1% deoxycholate, 1 mM EDTA, 500 mM NaCl and TE buffer. Finally, using elution buffer (50 mM Tris–HCl, 1 mM EDTA, 1% SDS) incubated at 65°C for 10 min, protein/DNA complexes were detached from beads. After removal of magnetic beads, complexes were de-crosslinked overnight at 65°C. Following treatment with 30ug RNaseA (30 min at 37°C) and 120ug proteinase K (2–3 hr at 37°C), DNA was purified using the Qiagen MinElute PCR Purification Kit using manufacturer’s instructions.

### Sequencing

#### Analysis of published single cell sequencing dataset

Raw data files were downloaded from GEO Accession: GSE146811. The data were processed using Seurat Package following the pipeline described by the Satija Lab: https://satijalab.r-universe.dev. Following SCTransformation to normalize data, we subset the data into only the first 5 timepoints (T00, T01, T02, T03 and T04) corresponding to Intact, 1-day, 7-day 14-day and 28-day regressed anterior prostate lobes. We then subset the data to include only clusters positive for EpCam, corresponding to all epithelial populations in the prostate (basal and luminal cells in all states). All Dot plots represent the expression by timepoint of a given gene within this entire epithelial subset.

#### ChIP sequencing analysis

Reads were trimmed using Trimmomatic (v0.6.6) using default parameters and then aligned to the mm10 genome using BWA (v0.7.17). Removal of PCR duplicates was then done using Picard (v2.0.1). MACS2 was then used for peak identification using default parameters and inputs as controls. To determine the significance of detected peaks, MACS2 used a Poisson distribution model to identify peaks and were corrected for false discovery rate using Benjamini-Hochberg correction. Aligned BAM files were then used to generate the bigwigs using DeepTools. Following annotation of all peaks using the ChIPSeeker R package, gene lists were generated from all genes with promoter peaks for further comparison, and Gene Ontology analysis was done using the clusterProfiler R package.

## Table of Antibodies

| Target | Species | Company & Cat# | Dilution |
| --- | --- | --- | --- |
| E-Cadherin | Rat | Invitrogen #13-1900 | 1:200 IF |
| Krt8 | Guinea pig | Fitzgerald #20R-CP004 | 1:200 IF |
| Krt5 | Chicken | BioLegend #905901 | 1:300 IF |
| cl-caspase 3 | Rabbit | Cell signalling #9661 | 1:100 IF |
| LAMP-1 | Mouse | Abcam #ab24170 | 1:100 IF |
| CD68 | Mouse | Invitrogen #MAF-13324 | 1:200 IF |
| PanKLa | Rabbit | PTM Bio # PTM-1401RM | 1:100 IF; 1:500 WB |
| Histone-H3 | Rabbit | Abcam #ab1791 | 1:10 000 |
| Tacstd2 | Goat | R&D Systems BAF1122 | 1:100 IF |

## Author Contributions

A.L.G.P. conceived the project. A.L.G.P. and L.M. directed the work, designed experiments, and interpreted data A.L.G.P. performed the experiments. A.L.G.P., S.V. and M.T. generated mice used for experiments and performed mouse surgeries. M.W., W.A.P, and D.S. supported data analysis. The manuscript was written by A.L.G.P and edited by L.M, M.T., W.A.P., and D.S. All authors discussed the data and contributed to the manuscript.

