## Supplemental Figures for "Engulfing to Adapt: Efferocytosis by Epithelial Cells Triggers Cell State Transitions During Tissue Remodeling"

Supplemental Figure 1

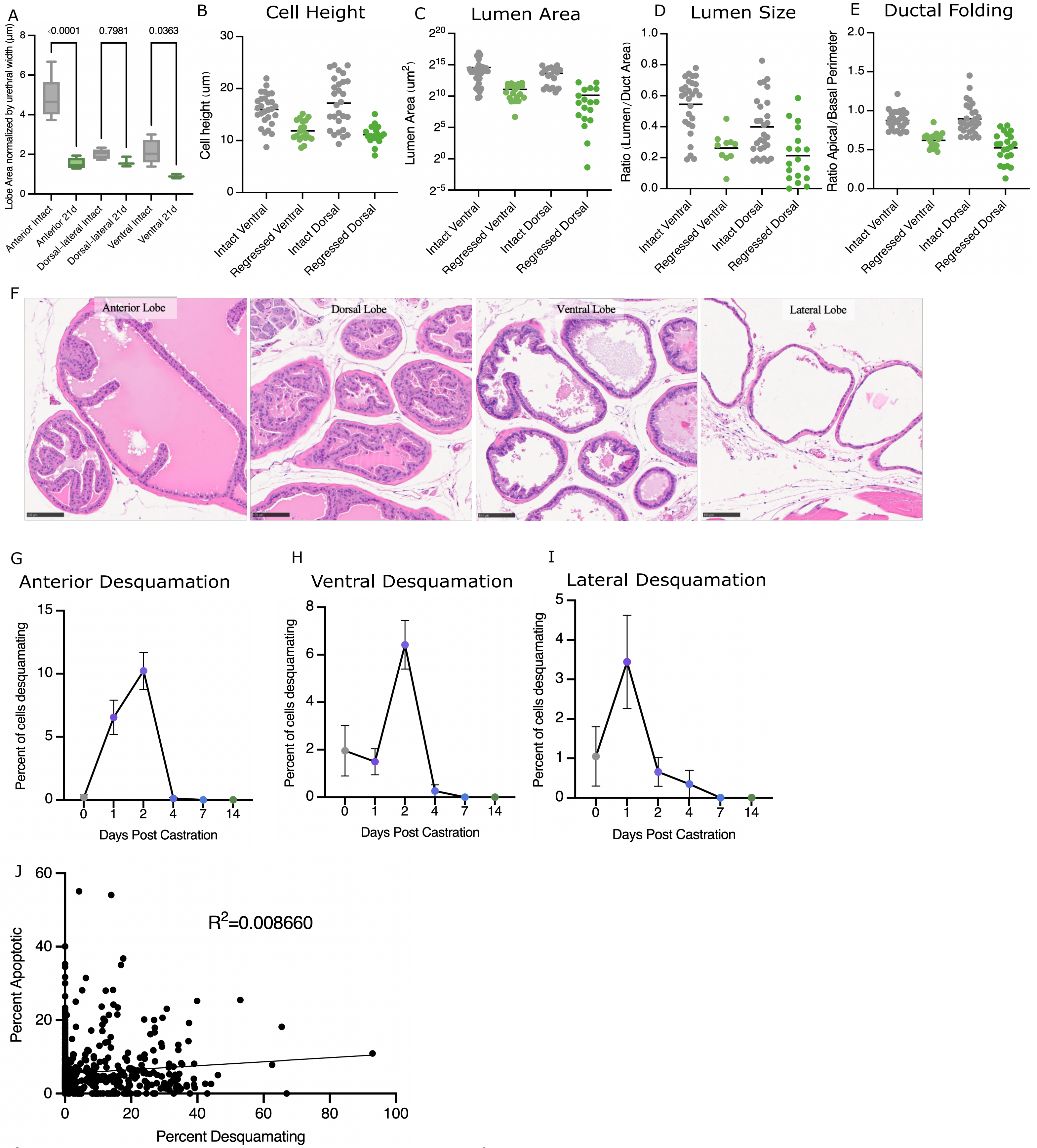


**Supplementary Figure 1. Morphological regression of the mouse prostate is due to desquamation, apoptosis and changes to cell shape.**

A. Lobe area measurements, in um2, each normalized by urethral width. N=3 mice.

B-E are measurements taken from the dorsal and ventral lobes, N=3 mice, 10 ducts per mouse per lobe.

B. Cell height, each data point represents average cell height per duct.

C. Area of lumen, each data point represents the lumen of one duct.

D. Lumen size, quantified as ratio of lumen area/ ductal area to control for changing duct size with regression, each data point represents one duct.

E. Ductal folding is a ratio of apical to basal ductal perimeters, each data point represents one duct.

F. Representative H&E images of anterior, dorsal, ventral and lateral lobes of an intact prostate to show the difference in morphology.

G. Average percent of cells in the duct found to be detached in the anterior lobe over time, Error bars= SEM.

H. Average percent of cells in the duct found to be detached in the ventral lobe over time, Error bars= SEM.

I. Average percent of cells in the duct found to be detached in the lateral lobe over time, Error bars= SEM.

J. Percent apoptotic cells versus percent desquamating cells per duct from 1-2 days regression. Each dot represents a single duct. All lobes are merged. n=4 mice, 20-50 ducts per mouse per lobe. Linear regression slope of 0.02265 with an R2=0.008660.

Supplemental Figure 2

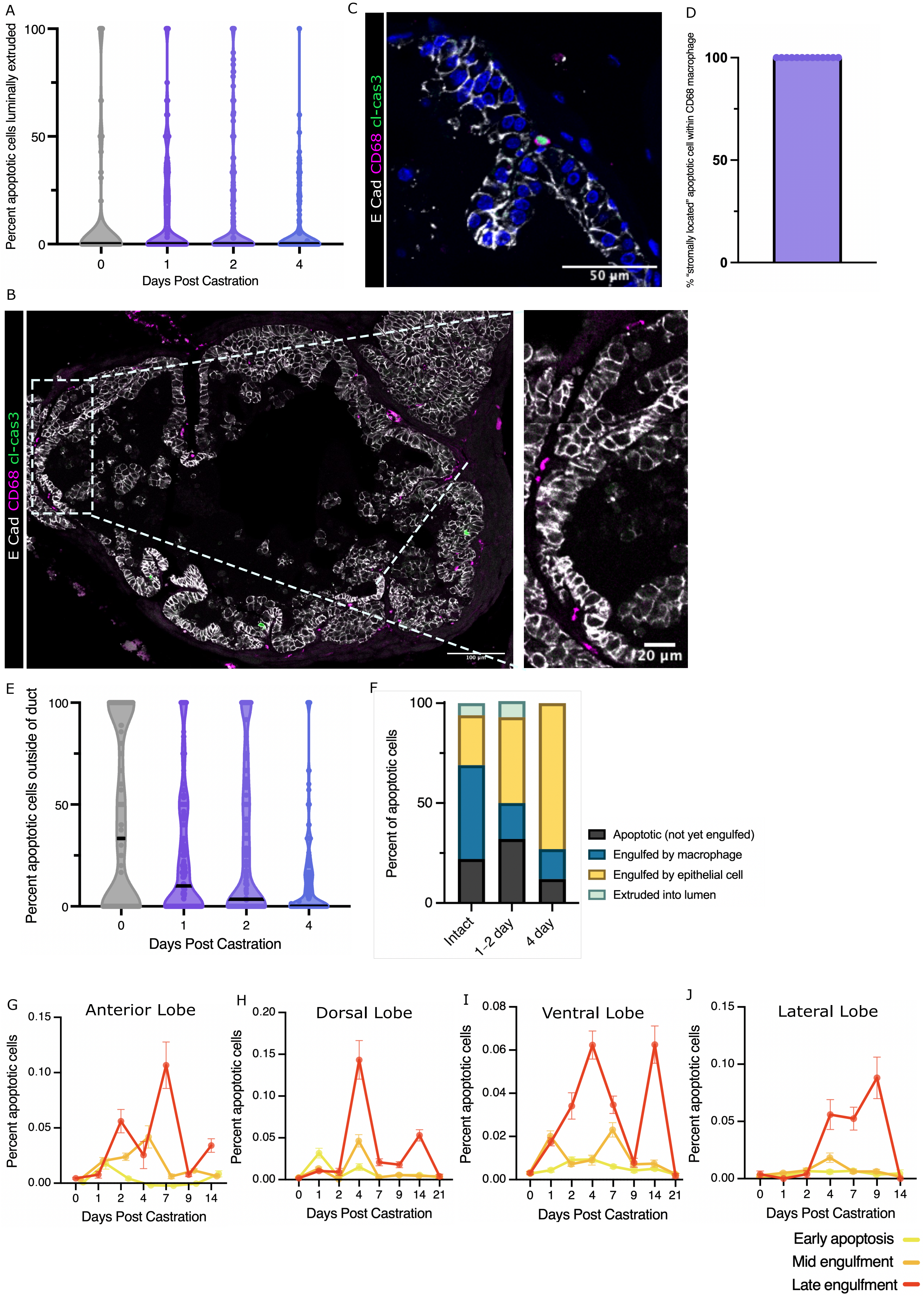

**Supplementary Figure 2. Epithelial cells in the prostate are phagocytic in response to castration-induced apoptosis**

A. Violin plots showing the percent of apoptotic cells that are found extruded in the lumen. Each data point represents the percentage within a given duct. Solid black line represents the median. Dotted lines show the other quartiles. Data compiled from 3 mice per timepoint, from 50-75 ducts per mouse. Mean values (not shown on graph) are as follows: intact 10%, 1d 14%, 2d 14% and 4d 6%.

B. Tissue section of a prostate duct 1 day after castration with immunofluorescence stains for E-cadherin, cleaved caspase 3 to label apoptotic cells and CD68 to show macrophages associated with the duct.

C. Immunofluorescence-stained tissue section of prostate duct 1 day after castration to show the localization of a cl-cas3 positive body within a CD68 macrophage located just basally outside of the duct.

D. Quantification of the percent of “stromally located” cl-cas3+ bodies, those located just outside of the duct on the basal side, that can be found within a CD68+ cell.

E. Violin plots showing the decrease in the contribution of macrophage efferocytosis in early regression compared to intact prostates. Each data point represents the percentage of total apoptotic cells in or around a duct that can be found within a macrophage. Solid black line represents the median. Dotted lines show the other quartiles. Data compiled from 3 mice per timepoint, from 50-75 ducts per mouse. Mean values (not shown on graph) are as follows: intact 41.8%, 1d 23.85%, 2d 28.66% and 4d 12.85%.

F. Distribution of apoptotic cells in and around epithelium. For each time point, representing intact, early (1-2d) and mid-regression (4d) the average (mean) was taken of the percentage of apoptotic cells found to be (A black) apoptotic but not yet engulfed or extruded, (B yellow) engulfed by epithelial cell, (C blue) engulfed by macrophage or (D green) luminally extruded. The SEM (not plotted for clarity) for each mean value is as follows: Intact A=5.75, B=5.51, C=7.23, D=3.21; 1d-2d A=3.37, B=3.3, C=2.87, D=2.04; 4d. A=2.25, B=4.03, C=2.41, D=0.19.

G-I. Of the apoptotic cells present in a duct, the average percent that is either in the early stages of being engulfed by an epithelial cell (yellow), mid-engulfed (orange) or late stage of engulfment (red) are plotted throughout regression. Data averaged from 3 mice per timepoint, from 8-10 ducts per timepoint per lobe. Error bars represent SEM.

Supplemental Figure 3

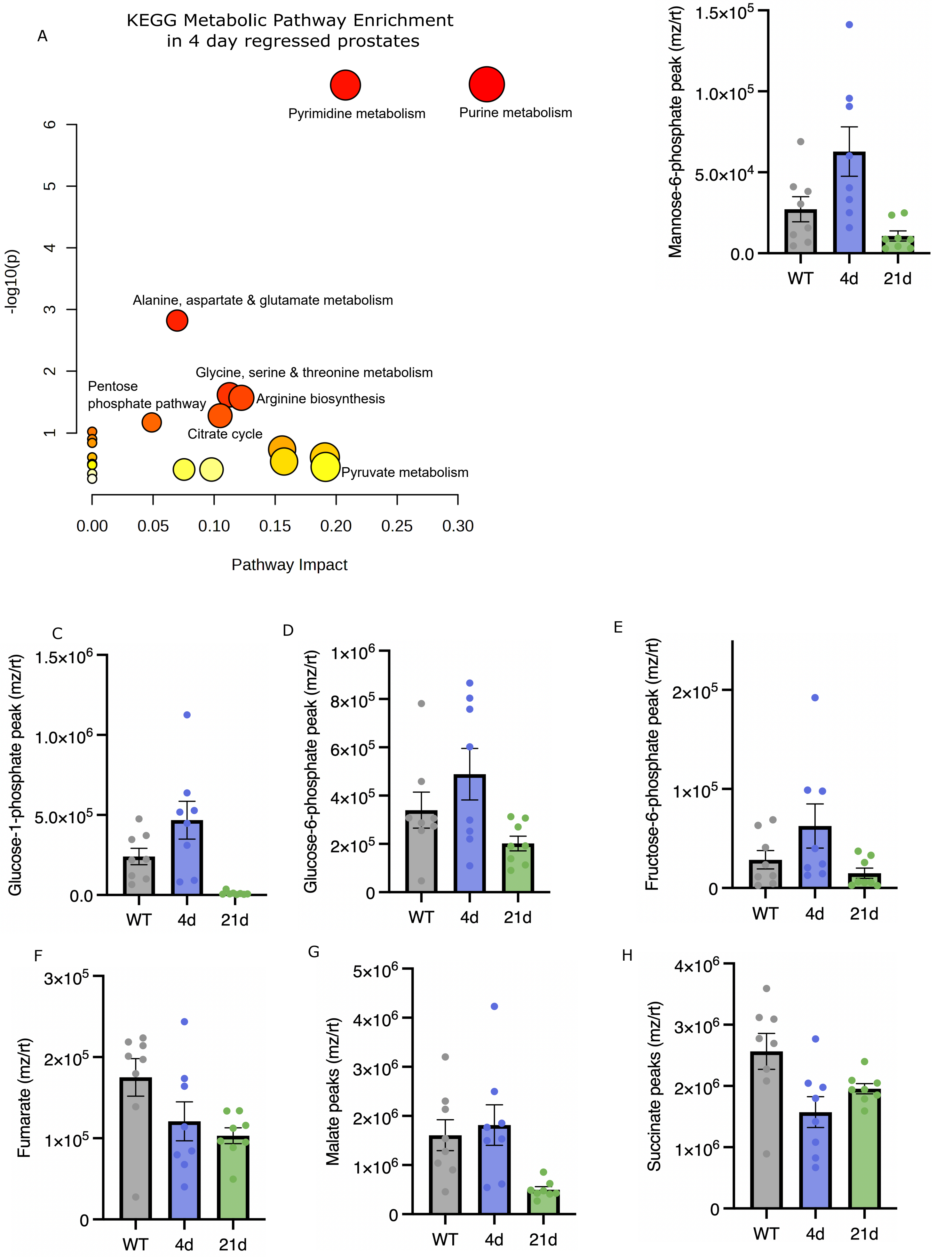

**Supplementary Figure 3. Prostate epithelial cells adapt their metabolism to accommodate efferocytic demand**
A. Metabolic pathways enriched, according to metabolites significantly ($p<0.05$) increased in 4-day regressed prostates versus intact.
B-H. Levels of metabolites present in anterior & dorsal lobes of intact, 4- and 21- regressed prostates. Each data point represents the metabolite level per mouse, 8 mice per condition. Error bars represent SEM. ANOVA statistical tests for differences in B. Mannose-6-phosphate between Intact_4d p=0.0511, 4d_21d p=0.0039, Intact_21d p=0.4895. C. Glucose-1-phosphate between Intact_4d p=0.1030, 4d_21d p=0.0008, Intact_21d p=0.0983. D. Glucose-6-phosphate between Intact_4d p=0.3749, 4d_21d p=0.0396, Intact_21d p=0.4298. E. Fructose-6-phosphate between Intact_4d p=0.2316, 4d_21d p=0.0684, Intact_21d p=0.7810. F. Fumarate between Intact_4d p=0.1598, 4d_21d p=0.8088, Intact_21d p=0.0484. G. Malate between Intact_4d p=0.8762, 4d_21d p=0.0147, Intact_21d p=0.0426. H. Succinate between Intact_4d p=0.0151, 4d_21d p=0.4733, Intact_21d p=0.1663.

Supplemental Figure 4

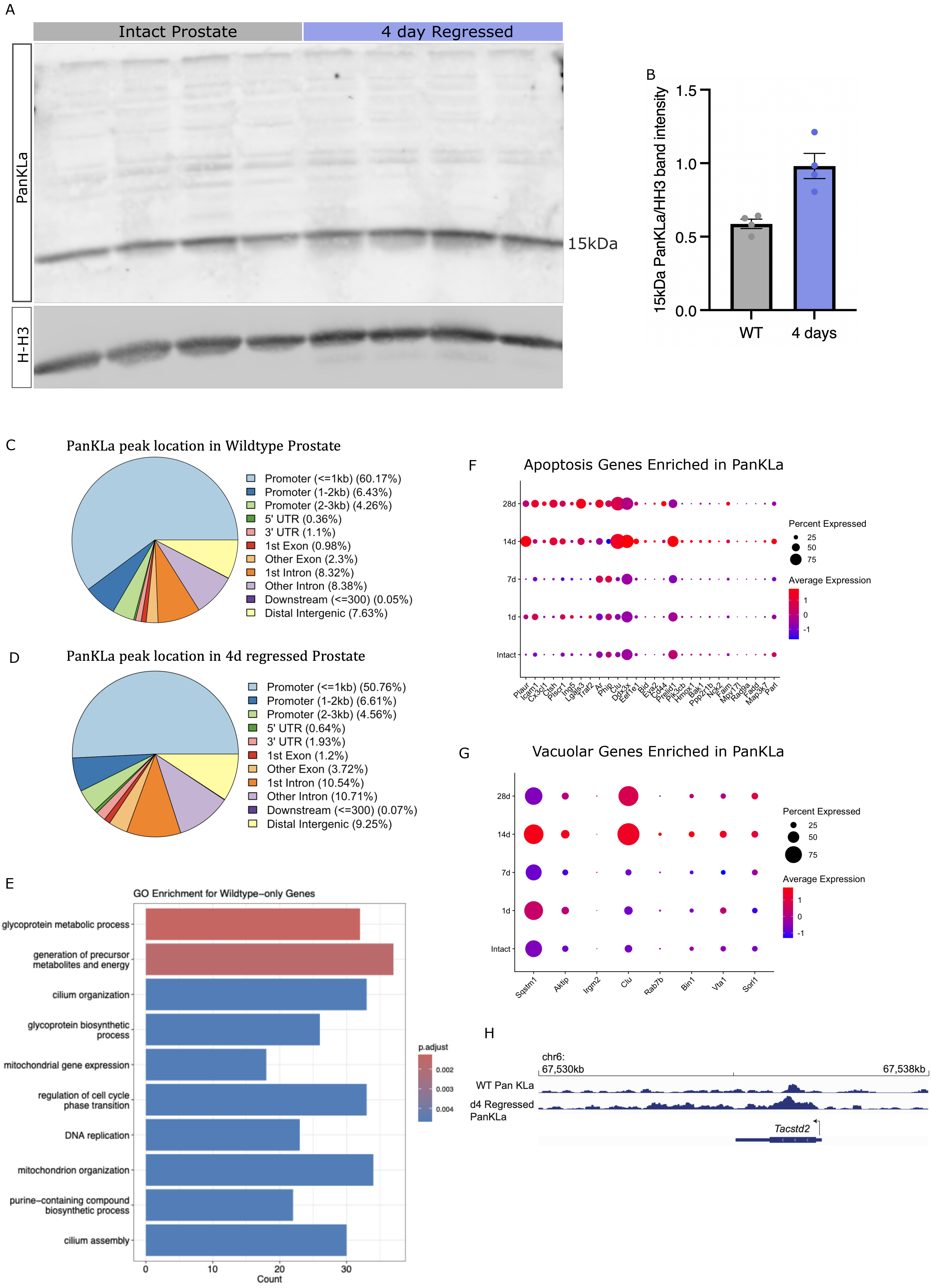

**Supplementary Figure 4. Histone lactylation marks in early regression align with genes important for efferocytosis and autophagy**

A. Western blot of intact and 4-day regressed prostate protein lysates (each lane represents a different mouse). Blotted for PanKLa and Histone H3.
A-B. Quantification of 15kDa PanKLa intensity, normalized by H-H3 intensity.
C-D. Distribution of PanKLa peaks across the genome in intact and 4-day regressed prostates.
E. GO terms of enriched pathways for genes with promoter lactylation exclusively in intact prostates.
F. Dot plot of single cell gene expression of apoptosis regulation genes with promoter lactylation exclusively in 4-day regressed prostates throughout regression, in epithelial cells.
G. Dot plot of single cell gene expression of vacuole-related genes with promoter lactylation exclusively in 4-day regressed prostates throughout regression, in epithelial cells.
H. Example track of promoter lactylation (PanKLa ChIP seq) upstream (and exonic) of the gene Tacstd2.

Supplemental Figure 5

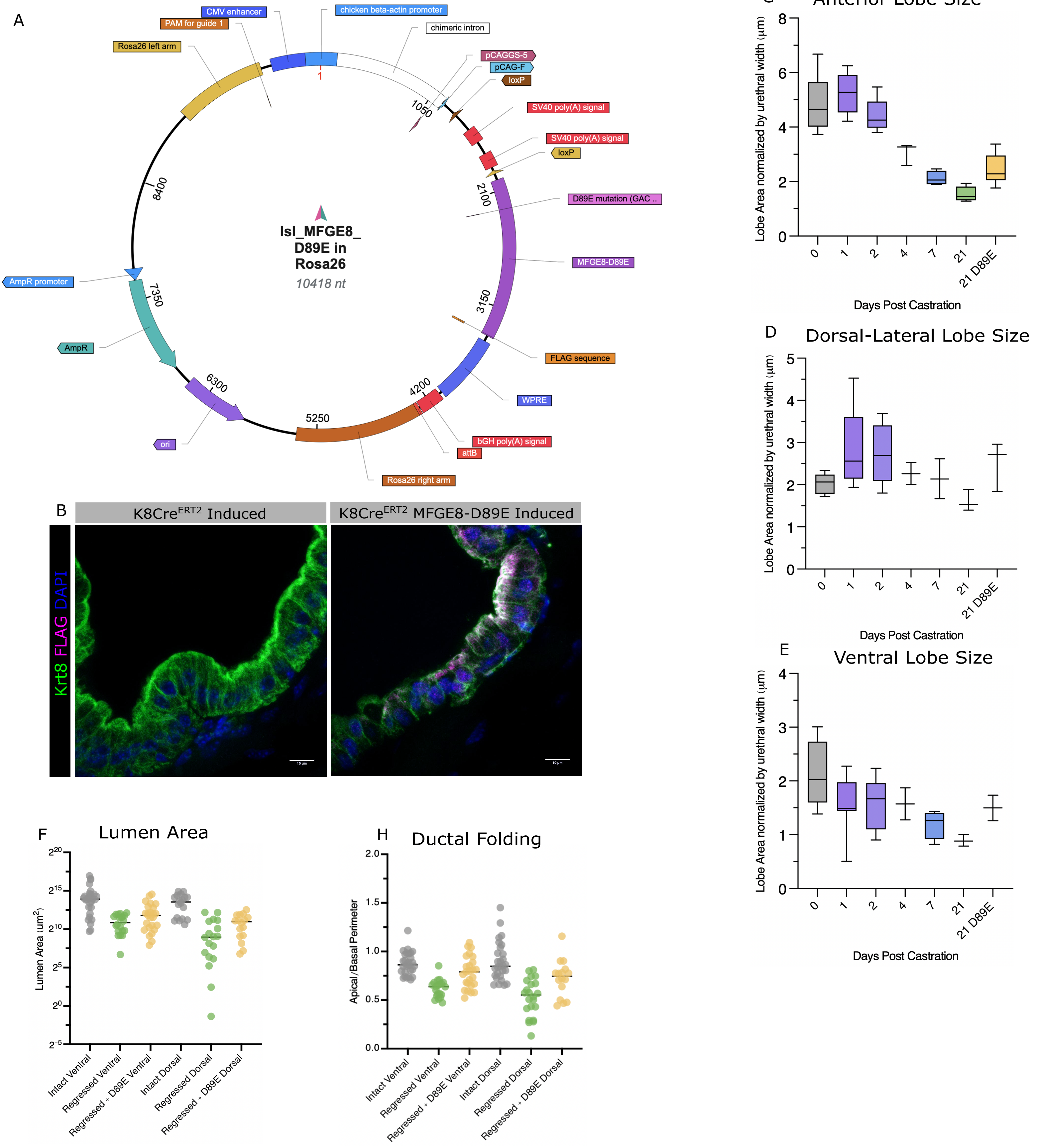


**Supplementary Figure 5. Impairing efferocytosis impedes prostate regression**
A. Vector image of MFGE8-D89E vector used to produce mouse, generated using VectorBee software.
B. Tissue section of a prostate duct 1 week after tamoxifen induction of MFGE8-D89E expression with immunofluorescence stains for Krt8 to label luminal cells, DAPI and FLAG to detect MFGE8-D89E protein expression.
C. Anterior lobe size, measured from 2D whole mount images of the frontal-oriented prostate and normalized to urethral width. Each boxplot represents the distribution of 3-5 mice per timepoint.
D. Dorsal-Lateral lobe size, measured from 2D whole mount images of the sagittal-oriented prostate and normalized to urethral width. Each boxplot represents the distribution of 2-3 mice per timepoint.
E. Ventral lobe size, measured from 2D whole mount images of the sagittal-oriented prostate and normalized to urethral width. Each boxplot represents the distribution of 2-3 mice per timepoint.
F. Area of lumen, each data point represents the lumen of one duct. N=3 mice, 10 ducts per mouse per lobe.
G. Ductal folding is a ratio of apical to basal ductal perimeters, each data point represents one duct. N=3 mice, 10 ducts per mouse per lobe.

Supplemental Figure 6

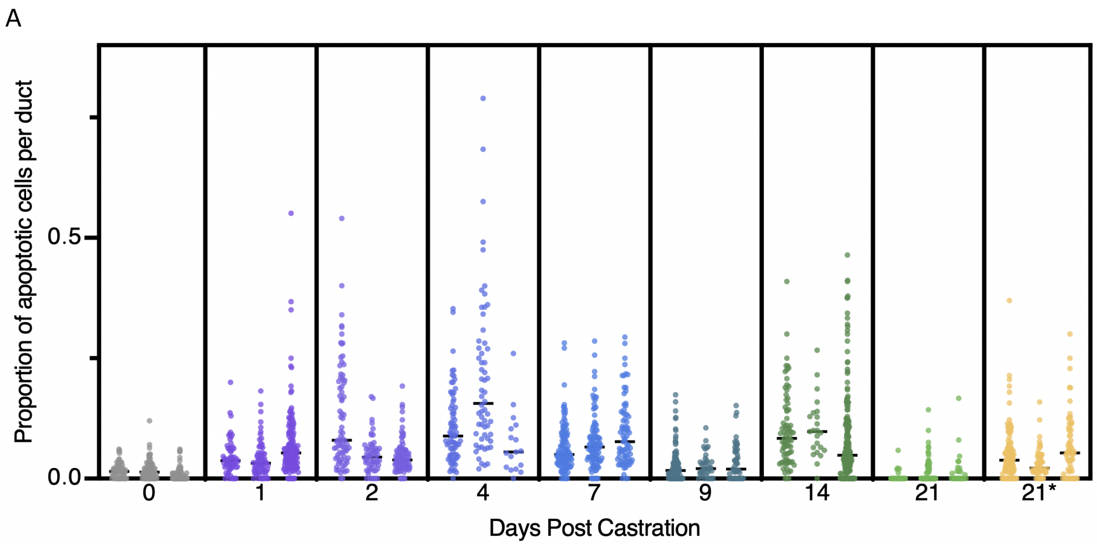


**Supplementary Figure 6. Proportion of apoptotic cells in prostate ducts after castration**
A. Scatter plot of the proportion of cells that are apoptotic, with individual mouse replicates separated to show variability. Each data point represents the percent apoptotic (TUNEL+) nuclei per duct, compiled from 3 mice per timepoint, from 20-75 ducts per mouse.
